# pH-dependent beta-lactam resistance in *Klebsiella pneumoniae* is mediated by paralogous class B PBPs and the class A PBP, PBP1b

**DOI:** 10.1101/2025.03.12.642896

**Authors:** Sarah Beagle, Petra Anne Levin

**Affiliations:** Washington University in St. Louis, Department of Biology, St. Louis, Missouri, USA

## Abstract

*Klebsiella pneumoniae* is a leading cause of global deaths due to antibiotic resistance. Of particular concern is the rapid expansion of resistance to beta-lactam antibiotics within *K. pneumoniae* lineages. The environmental factors that influence pathogen physiology and, subsequently, antibiotic resistance remain poorly understood. Here we demonstrate that physiologically-relevant reductions in pH increased *K. pneumoniae* beta-lactam resistance as much as 64-fold, with the most dramatic increase observed for beta-lactams that specifically inhibit cell division. We identified two genes that contribute to acid-dependent beta-lactam resistance, the class A PBP, PBP1b, and the paralogous class B PBP, PBP3_PARA_. Loss of either PBP1b or PBP3_PARA_ increases *K. pneumoniae* susceptibility to beta-lactams at low pH. Altogether these data emphasize the importance of functional redundancy among cell wall synthesis enzymes which allows for specialization and ensures robust cell wall synthesis across a range of environmental conditions.

**Importance:** Beta-lactams are the most prescribed class of antibiotics, but their effectiveness is threatened by a global rise in antimicrobial resistance. How the environment within a host or infection site shapes pathogen response to antibiotics is frequently overlooked in assessments of antibiotic effectiveness. We demonstrate that growth at physiologically-relevant low pH substantially increases *Klebsiella pneumoniae* resistance to clinically important beta-lactams. An important finding of this study is that during growth in acidic pH *K. pneumoniae* has a different repertoire of cell wall synthesis genes available than during growth at neutral pH due to the presence of acid-inducible paralogous copies of essential cell wall synthesis enzymes, PBP2 and PBP3. An additional functionally-redundant enzyme, PBP1b, also contributes to acid-dependent beta-lactam resistance. Together, these findings expand our understanding of how bacteria maintain cell wall synthesis across diverse physiochemical environments and highlight potential new therapeutic targets.

## Introduction

Infections caused by the ESKAPE pathogen *K. pneumoniae* are increasingly difficult to treat due to dramatic rises in antibiotic resistance. *K. pneumoniae* is a dominant member of two different Beta-lactam-resistant groups of bacteria, <u>E</u>xtended-<u>s</u>pectrum-<u>B</u>eta-lactamase (ESBL)-producing *Enterobacteriaceae* and <u>C</u>arbapenem-resistant *<u>E</u>nterobacteriaceae* (CRE) which are labeled serious and urgent public health threats by the CDC^1^. In 2019, *K. pneumoniae* was the third leading cause of global deaths where the death was directly due to or associated with antibiotic resistance^2^. While ESBLs and related enzymes are major contributors to beta-lactam resistance in Gram-negative bacteria, data from our group and others identifies physicochemical parameters consistent with a host environment are under-appreciated drivers of resistance. Changes in pH, nutrient availability, and accumulation of the alarmone, ppGpp, increase antibiotic resistance across a range of bacterial pathogens, highlighting the need to better understand how environmental factors impact pathogen physiology^3–6^.

Despite the prevalence of resistance markers such as ESBLs, beta-lactams remain the most prescribed antibiotic class^7^. Beta-lactams interfere with the synthesis of the cell wall or peptidoglycan (PG), an essential structure that provides shape and structural rigidity. PG is comprised of glycan strands of repeating linked N-acetylglucosamine (NAG) and N-acetylmuramic acid (NAM) sugars that are crosslinked together by short peptide chains ^8^. Synthesis and maintenance of PG requires many enzymatic steps that occur across multiple cellular locations; PG precursors are built in the cytoplasm, transported through the inner membrane, and fully assembled into PG within the periplasmic space. Amongst the *Enterobacteriaceae,* periplasmic PG assembly is highly enzymatically redundant, with an average of four enzymes capable of carrying out a single periplasmic PG synthesis reaction^9^. This apparent functional redundancy is particularly striking when compared to the earlier cytoplasmic PG synthesis reactions, which occur in a nearly 1:1 enzyme-to-reaction ratio^10^. While precursors are assembled in the relatively buffered cytoplasm, the cell wall itself is built in the periplasm and thus exposed to the physicochemical environment.

Penicillin-binding proteins (PBPs) carry out essential transpeptidation reactions in the periplasm. Class A PBPs are bi-functional enzymes that both link the NAG-NAM sugar moieties together (transglycosylation, TG) as well as crosslink the peptide stems of adjacent glycan strands (transpeptidation, TP). In *Escherichia coli,* the primary class A PBPs are PBP1a and PBP1b, which are encoded by *mrcA* and *mrcB* respectively. PBP1a and PBP1b are individually dispensable for growth, but loss of both is lethal^11–13^. A third class A PBP, PBP1c (*pbpC*), has an unknown role in PG synthesis and maintenance and is dispensable under standard laboratory conditions^14^. Class B PBPs are monofunctional transpeptidases; these PBPs work in conjunction with cognate glycosyltransferases. PBP2 and PBP3 are the canonical class B PBPs in Gram-negative bacteria, and they function within the elongation and division machinery respectively.

Beta-lactams interfere with PG synthesis by irreversibly binding to the transpeptidase domain of PBPs and inhibiting transpeptidation^7^. Our previous work indicates that growth in acidic pH (∼pH 5) substantially increases *E. coli* resistance to PBP2- and PBP3-targeting Beta-lactams. Both growth in pH values < 5, as well as acid-dependent Beta-lactam resistance required the Class A PBP, PBP1b, while full fitness during growth in alkaline pH required PBP1a^3^. To determine if this response is conserved amongst other *Enterobacteriaceae*, we evaluated the impact of environmental pH on *K. pneumoniae* resistance to beta-lactams.

Here we report that *K. pneumoniae* beta-lactam resistance increased as much as 64-fold during growth in acidic conditions. Genetic analysis identified two cell wall synthesis genes, *mrcB* and *ftsI2,* that are critical for pH-dependent changes in antibiotic resistance. *mrcB* encodes PBP1b, a bifunctional class A PBP previously identified as a contributor to pH-dependent antibiotic resistance in *E. coli*^3^. Induced at low pH, *ftsI2* encodes PBP3_PARA,_ a paralogous copy of the canonical, division-associated class B PBP, PBP3. PBP3_PARA_ is required for both acid-dependent beta-lactam resistance and robust growth and division at low pH. Altogether our findings highlight that functional redundancy in bacterial PBPs confers fitness advantages and contributes to environmentally-driven antibiotic resistance.

## Results

### *Klebsiella pneumoniae* exhibits acid-dependent resistance to Beta-lactam antibiotics

To determine the impact of pH on *K. pneumoniae* beta-lactam susceptibility, we assessed the response of a urinary tract infection (UTI) isolate, *K. pneumoniae* TOP52^15^ to a panel of beta-lactams during growth across a range of physiologically relevant pH conditions. The drug panel included beta-lactams that preferentially target PBPs in the divisome (i.e. Cephalexin, Aztreonam, and Piperacillin) or the elongasome (Meropenem, Doripenem), as well as non-specific inhibitors (Ampicillin, Carbenicillin) that are reported to have similar affinities for all PBPs^16,17^. Relative susceptibility or resistance to a given Beta-lactam was determined by comparing the minimal inhibitory concentration of drug at neutral pH to that at other pH values.

*K. pneumoniae* exhibited a 2-to 64-fold increase in resistance to the entire spectrum of tested beta-lactams when grown in LB at pH values < 5 relative to resistance at pH 7. (Fig 1A, Supp Table 4). We observed strong acid-dependent increases in resistance for drugs that preferentially target PBP3, with Aztreonam (AZT) and Cephalexin (CEX) MICs increasing as much as 20- and 64-fold respectively at pH 4.8. Resistance to PBP2-targeting Beta-lactams increased modestly at low pH, with MICs to Doripenem (DOR) and Meropenem (MER) increasing 2-to 4-fold. In contrast to *E. coli*^3^, we also observed modest 2-to 4-fold increases in resistance to the non-specific Beta-lactams, Ampicillin (AMP) and Carbenicillin (CAR) at low pH. At these pH values, increases in MICs during growth at low pH are not due to significant antibiotic degradation^3^.

**Fig 1.**
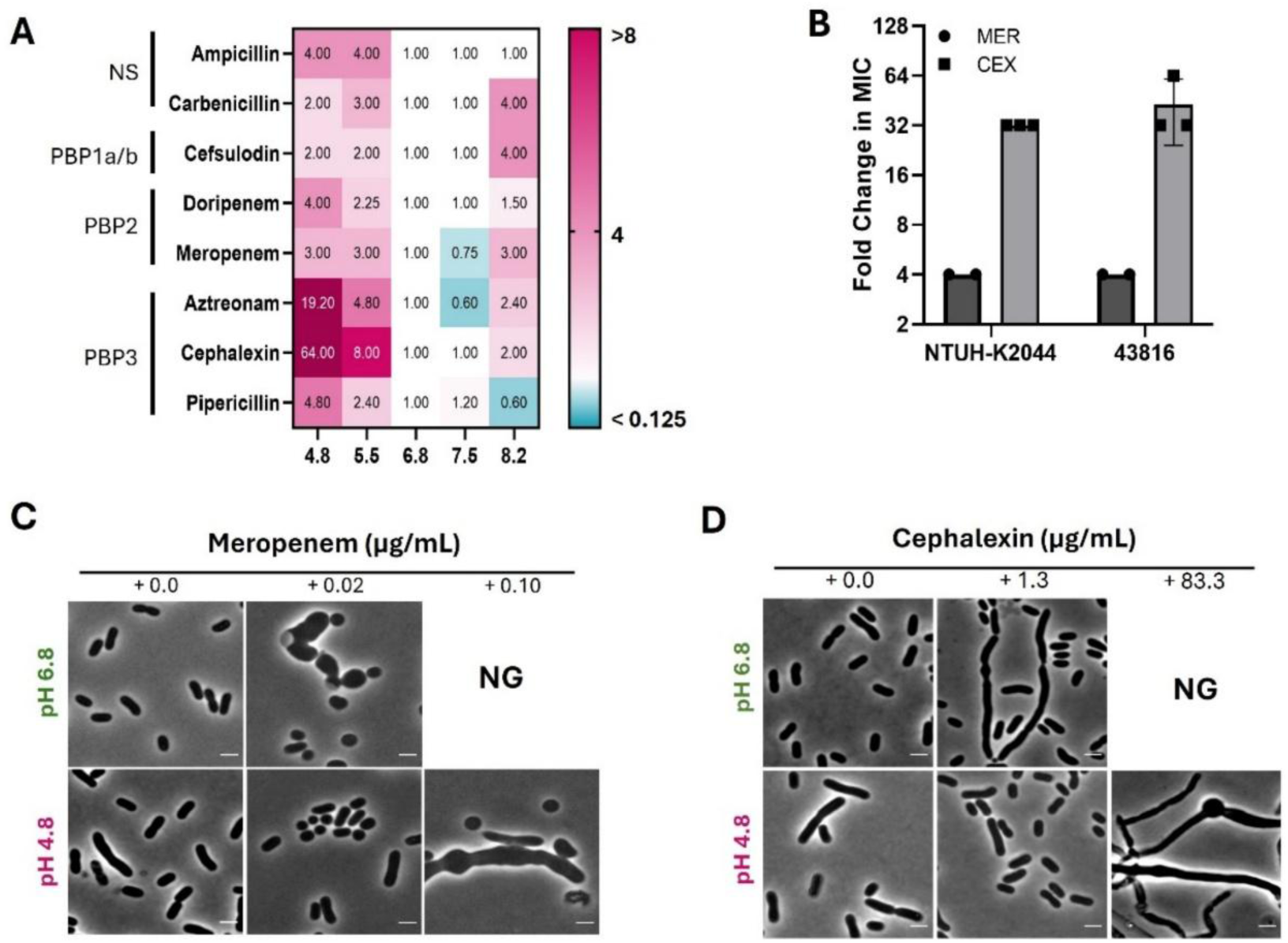
*Klebsiella pneumoniae* exhibits acid-dependent resistance to Beta-lactams. (A) Heatmap displaying the fold-change in minimal inhibitory change (MIC) for *K. pneumoniae* TOP52 at indicated pH relative to neutral pH. Each cell represents the median fold-change value from a minimum of three biological replicates performed in LB buffered to indicated pH with MMT buffer. Specific Beta-lactam antibiotics along with their preferential PBP target are indicated on the Y-axis. NS = non-specific (B) The fold-change in Meropenem (MER) or Cephalexin (CEX) MIC of *K. pnuemoniae* strains ATCC 43816 and NHUT-K2044 grown at pH 4.8 compared to growth at pH 6.8. Bars represent mean and error bars denote standard deviation. C-D) Representative micrographs of terminal morphologies of *K. pneumoniae* TOP52 following growth for 20 hours in LB (pH 4.8 or 6.8) and treated with Meropenem (C) or Cephalexin (D) at indicated concentrations. Scale bar = 2 µm.

Low pH not only enabled growth at concentrations of drug that are lethal at neutral pH but also preserved cell morphology (Fig 1 C & D). Consistent with increased intrinsic resistance at low pH, the characteristic rod-shaped morphology of *K. pneumoniae* was preserved during growth under acidic conditions in concentrations of beta-lactams that cause either cell rounding (Meropenem) or filamentation (Cephalexin) at neutral pH. The observed acid-dependent increased resistance to beta-lactams is not strain specific behavior as *K. pneumoniae* strains NTUH-K2044 and ATCC 43816 also exhibit a 4-fold increase to Meropenem (MER) and an ∼32-fold increase to Cephalexin (CEX) when grown at pH 4.8 versus pH 6.8 (Fig. 1B).

To determine if this behavior also occurs under conditions that better reflect a native host environment, we assayed how *K. pneumoniae* responded to beta-lactams during growth in an artificial urine growth medium at low and neutral pH^18^. Similar to growth in LB, we observed 32-to >64-fold increases in resistance to Ampicillin and Cephalexin in acidified artificial urine with and without amino acid supplementation compared to neutral pH (Sfig 1, Supp table 4 & 5). While the MIC for a given beta-lactam was generally higher during growth at low pH independent of media composition, we observed that for certain antibiotics the media composition coupled with acidification of the media significantly altered the susceptibility. The MIC for Ampicillin at low pH in LB was significantly lower (mean MIC = 104 ug/mL ± 37) than during growth in AU (mean MIC = 2500 ug/mL ±0.0), indicating an interplay between nutrient composition and pH. We were unable to determine the exact foldchange in MIC for Cephalexin, as *K. pneumoniae* grew in all tested concentrations of Cephalexin in both AU and AU+AA (Sfig 1, table 4 & 5). Together our data suggest that acid-dependent beta-lactam resistance occurs across a range of nutrient conditions and is conserved amongst *K. pneumoniae* lineages.

### Acid-dependent Beta-lactam resistance is independent of endogenous beta-lactamase expression

*K. pneumoniae* TOP52 encodes an endogenous class-A beta-lactamase, *bla_SHV-1_,* which confers increased resistance to the Penicillin subclass of Beta-lactams^7,15,19^. To determine if expression of *bla_SHV-1_* is influencing the resistance profiles at low pH, we generated a deletion strain (Δ*bla _SHV-1_*) and repeated the MIC assays under both acidic and neutral pH conditions. We observed increased sensitivity to the Penicillin sub-classes of Beta-lactams tested (AMP, CAR, PIP) in both pH environments (Sup Table 5). Consistent with the specificity of *bla_SHV-1_*, loss of *bla_SHV-1_* did not alter the MICs to non-Penicillin beta-lactams at either pH (Sup Table 5). However, the Δ*bla _SHV-1_* mutant still displayed increased resistance to all tested Beta-lactams at low pH relative to neutral pH (Fig 2 A-D, Supp table 5), demonstrating that acid-dependent resistance is independent of endogenous beta-lactamase expression.

**Fig 2.**
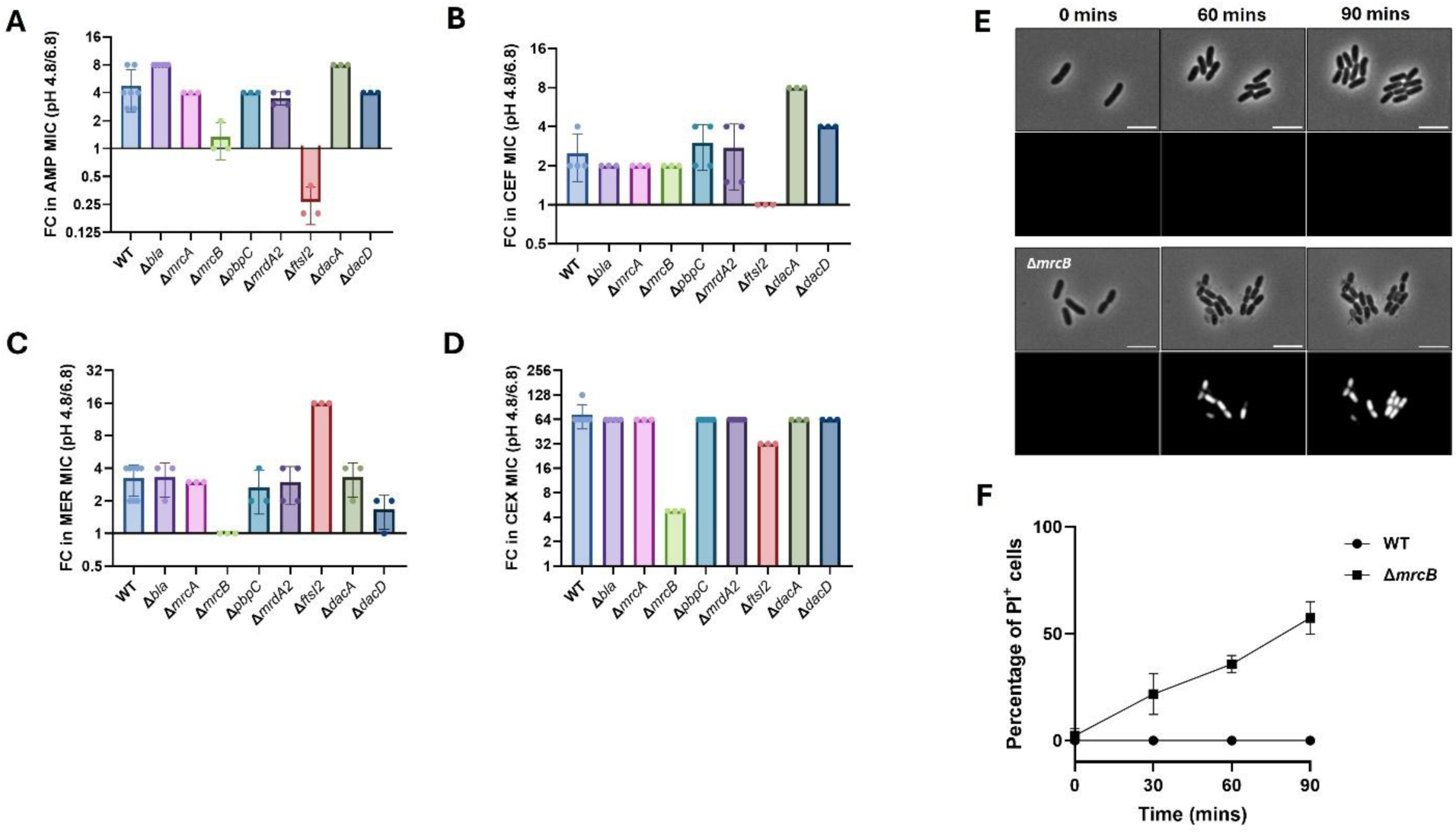
Loss of either PBP1b or PBP3_PARA_ results in differential impacts to beta-lactam resistance at low pH. (A-D) Fold change in MICs at pH 4.8 relative to pH 6.8 shown for generalist (A), class A-targeting (B), PBP2-targeting (C), and PBP3-targeting (D) beta-lactams. Data are graphed as median values with range. (E) Micrographs from time-lapse microscopy of WT (top) or Δ*mrcB* (bottom) growing in the presence of CEX at low pH. Microscopy pads comprised of LB buffered with MMT to pH 4.8 + 15.6 µg/mL CEX + 1.5 µM propidium iodide + 1% agarose. The CEX concentration reflects the 1X MIC for the Δ*mrcB* mutant and 0.06X MIC for the WT strain.

### pH modulates expression of Class B PBP paralogs in *K. pneumoniae*

To ascertain the impact of pH on expression of genes encoding PBPs or other cell wall synthesis genes, we performed RNAseq in *K. pneumoniae* comparing cells cultured in pH 4.8 LB versus pH 6.8 LB (Supp table 7). Consistent with previous studies, pH does not impact expression of class A PBPs, *mrcA* (PBP1a) or *mrcB* (PBP1b), in either *E. coli* or *K. pneumoniae*^20,21^ (Table 1). Class A PBP activity is regulated by cognate outer membrane activators, LpoA and LpoB^22–24^. We observed that the PBP1b activator, encoded by *lpoB,* was down-regulated ∼2-fold at low pH.PBP1c, a cryptic class A PBP with no demonstrated role in cell wall synthesis under standard laboratory conditions, exhibited an ∼2-fold decrease in expression during growth in acidic pH (Table 1, Supp table 7). We observed no significant changes expression of the class B PBPs between pH conditions, but there was an order of magnitude increase in expression at low pH of genes we designated *mrdA2* and *ftsI2*, which encode paralogous copies of class B PBPs, referred to here as PBP2_PARA_ and PBP3_PARA_ (Fig 3 A&B). In addition to the class B paralogs, several other cell wall synthesis genes displayed pH-responsive expression profiles. Most notably, two carboxypeptidases, *dacA* (PBP5) and *dacD* (PBP6b), exhibited modest increases of 2-to 3-fold in expression during growth at low pH. Both *dacA* and *dacD* are implicated in maintaining proper cell morphology in *E. coli*, with *dacD* being required for proper morphology specifically at low pH^25,26^.

**Fig 3.**
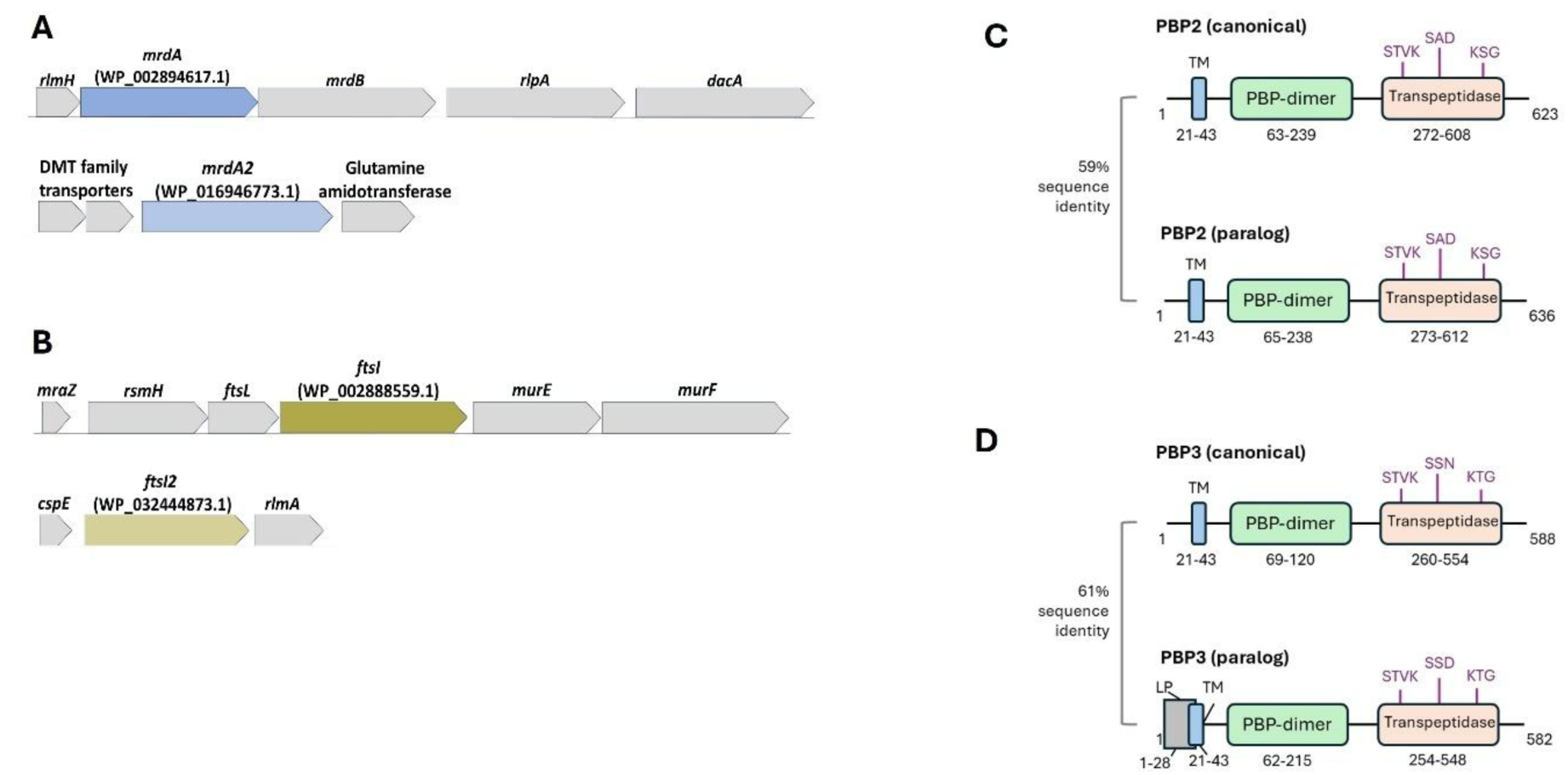
*Klebsiella pneumoniae* possesses acid-responsive paralogous copies of the Class B PBPs. (A) Genomic regions containing the canonical PBP2, encoded by *mrdA*, and the paralogous gene *mrdA2,* which encodes PBP2_PARA_ (B) Genomic regions containing the canonical PBP3, encoded by *ftsI,* and the paralogous gene *ftsI2,* which encodes PBP3_PARA_. (C & D) Domain conservation between canonical and paralogous copies of PBP2 (C) and PBP3 (D). Domains are denoted as follows: Transmembrane (TM), Lipoprotein lipid attachment site (LP, PS51257) PBP-dimer (Pfam: PF03717), Transpeptidase domain (Pfam: PF00905). The location of motifs that are critical for enzyme catalysis are highlighted in purple.

**Table 1.** Transcriptional response to pH of key beta-lactam targets in *K. pneumoniae* and *E. coli*.

|  | Gene | Enzyme Encoded | Description | log<sub>2</sub>FC<br>(pH 4.8/6.8) |
| --- | --- | --- | --- | --- |
| ***K. pneumoniae*** |  |  |  |  |
| D1637_RS05825 | *mrcA* | PBP1a | bifunctional glycosyl transferase/transpeptidase | **0.60** |
| D1637_RS12385 | *mrcB* | PBP1b | bifunctional glycosyl transferase/transpeptidase | **-0.39** |
| D1637_RS01040 | *pbpC* | PBP1c | peptidoglycan glycosyltransferase | **-1.61** |
| D1637_RS15400 | *mrdA* | PBP2 | canonical peptidoglycan DD-transpeptidase | **0.84** |
| D1637_RS12000 | *ftsI* | PBP3 | canonical peptidoglycan DD-transpeptidase | **0.23** |
| D1637_RS21590 | *mrdA2* | PBP2<sub>para</sub> | paralogous peptidoglycan DD-transpeptidase | **3.41** |
| D1637_RS24275 | *ftsI2* | PBP3<sub>para</sub> | paralogous peptidoglycan DD-transpeptidase | **3.28** |
| D1637_RS15385 | *dacA* | PBP5 | D-Ala-D-Ala-carboxypeptidase | **1.20** |
| D1637_RS25010 | *dacD* | PBP6b | D-Ala-D-Ala carboxypeptidase | **1.63** |
| D1637_RS17685 | *lpoB* | LpoB | Outer membrane activator of PBP1b | **-1.12** |
| ***E. coli*** |  |  |  |  |
| b3396 | *mrcA* | PBP1a | bifunctional glycosyl transferase/transpeptidase | **0.36** |
| b0149 | *mrcB* | PBP1b | bifunctional glycosyl transferase/transpeptidase | **-0.23** |
| b2519 | *pbpC* | PBP1c | peptidoglycan glycosyltransferase | **-0.43** |
| b0635 | *mrdA* | PBP2 | canonical peptidoglycan DD-transpeptidase | **0.47** |
| b0084 | *ftsI* | PBP3 | canonical peptidoglycan DD-transpeptidase | **-0.64** |
| b0632 | *dacA* | PBP5 | D-alanyl-D-alanine carboxypeptidase | **0.95** |
| b2010 | *dacD* | PBP6b | D-alanyl-D-alanine carboxypeptidase | **2.63** |
| B1105 | *LpoB* | LpoB | Outer membrane activator of PBP1b | **-1.33** |

\*Shaded genes represent those that were differentially regulated by pH as determined by our cutoff of |log<sub>2</sub>FC| > 1 and p < .05. Blue Log<sub>2</sub>FC values indicate that the gene was down-regulated during growth at pH 4.8 versus pH 6.8. Magenta Log<sub>2</sub>FC values indicate that the gene was up-regulated during growth at pH 4.8 versus pH 6.8.

### Loss of PBP3_PARA_ results in filamentation in *K. pneumoniae* at low pH

Duplications of the class B PBPs, PBP2 and PBP3, that have expanded the repertoire of cell wall synthesis enzymes have been described in several pathogens^27–29^. Class B PBP paralogs in *Salmonella enterica* serovar Typhimurium (*S.* Typhimurium) referred to as PBP2_SAL_ and PBP3_SAL_ are thought to be important for maximal fitness during growth at low pH. Expression of both PBP2_SAL_ and PBP3_SAL_ is maximally induced at low pH (<5.8), and loss of PBP2_SAL_ has been reported to impair rod-shape morphology under acidic environments^28,29^.

As with PBP2_SAL_ and PBP3_SAL_, PBP2_PARA_ and PBP3_PARA_ share significant sequence identity with their canonical counterparts (59% for PBP2_PARA_ and 61% for PBP3_PARA_) and retain the sequence motifs associated with catalysis, but their contributions to cell wall maintenance in *Klebsiella* are unknown (Fig 3 C&D, Sfig 2). We generated individual gene deletion strains and examined the mutants using cell size analysis and time-lapse microscopy to evaluate the role of the class B PBP paralogs in cell size and division. We cultured cells at both acidic and neutral pH and sampled cultures at early exponential phase (OD_600_ = 0.1-0.2) for cell size analysis. While loss of PBP2_PARA_ did not impact cell size or morphology at either pH, we found that during growth at pH 4.8 the PBP3_PARA_ deletion mutant exhibited heterogeneous cell lengths, with many cells displaying filamentous behavior indicative of cell division defects (Fig4 A&B, Sfig 3). Using a definition of filamentation as a cell length of >15 µM which represents ∼3X the average length of WT cells at pH 4.8 (3.9 ± 0.2 µM, Fig 4B), we determined that 21-58% of PBP3_PARA_ deficient cells within a given population were filamentous, with a smaller proportion of cells (3-17%) reaching lengths of >40 µM. The division defects associated with loss of PBP3_PARA_ were specific to growth in acidic conditions, as growth at neutral pH resulted in an average cell length for the PBP3_PARA_ deficient cells consistent with WT cell lengths (Fig 3C).

**Fig 4.**
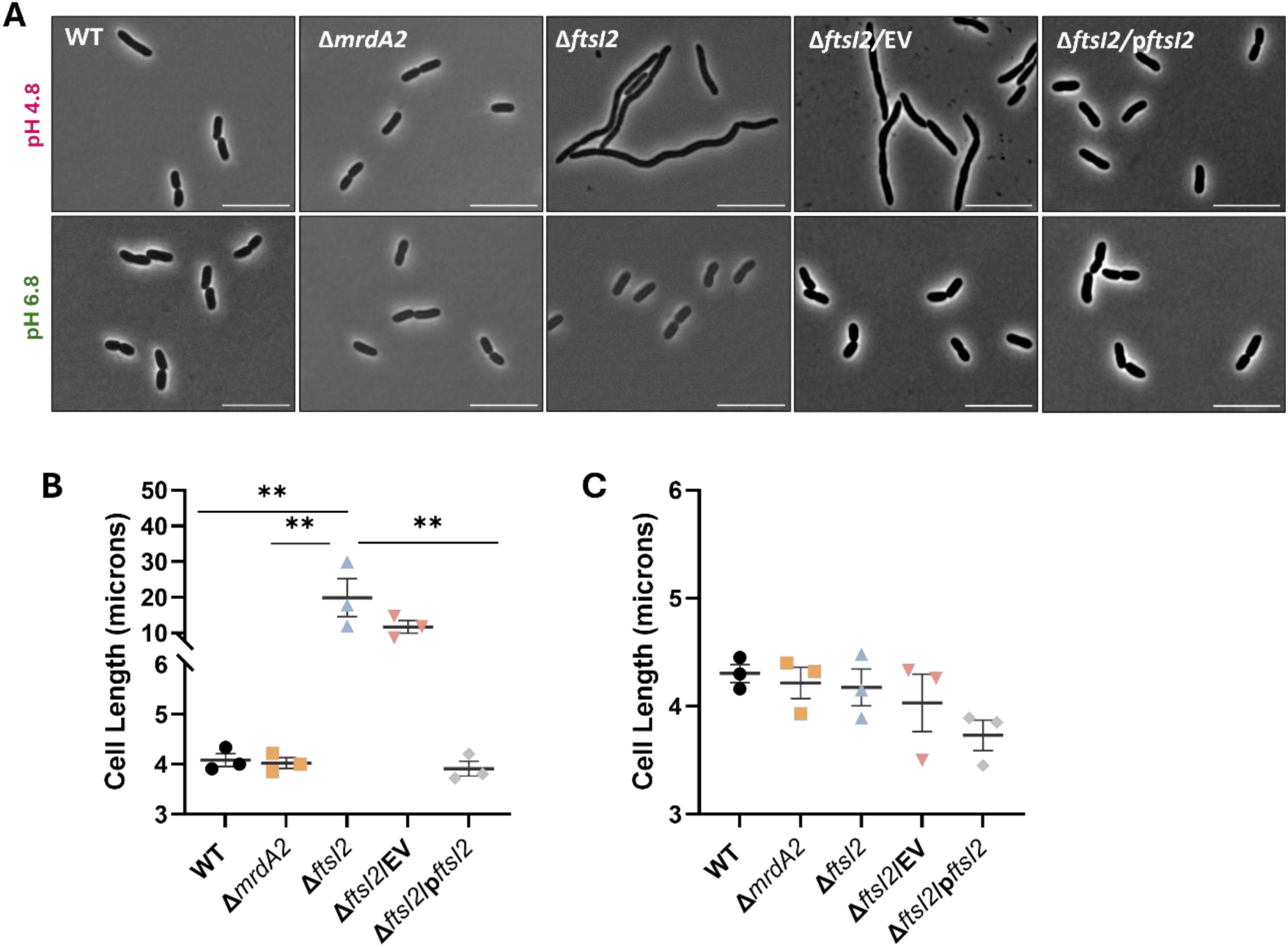
Loss of PBP3_PARA_ impairs division at low pH. (A) Representative micrographs of indicated strains following growth in LB buffered to pH 4.8 (top panels) or growth in LB buffered to pH 6.8 (bottom panels). For complementation studies with either empty vector (EV) or p*ftsI2,* the media was supplemented with 0.1 mM IPTG. Scale bar = 10 uM. (B-C) Quantification of cell length in (B). Each data point represents the mean of a biological replicate (>50 cells analyzed for each replicate). The bar represents the mean of three biological replicates, and the error bars represent the standard error of the mean. Statistical significance was determined by a one-way ANOVA with Tukey’s correction for multiple comparisons with asterisks denoting significance as follows:** = p<0.005, * = p<0.5.

Time-lapse microscopy of acid-grown Δ*ftsI2* cells spotted onto acidic LB agarose pads revealed a population of cells with heterogeneous lengths that displayed clear division defects as cells continued to elongate without successful division resulting in long filamentous cells. Whereas spotting acid-grown Δ*ftsI2* cells onto LB agarose pads at neutral pH restored division to the filamentous cells (Sfig 4). Complementation of *ftsI2* in trans from a medium-copy number plasmid enabled division and restored cell lengths consistent with WT at low pH (Fig 4 A&B). Together these data suggest that the activity of the canonical PBP3 is impaired under acidic conditions and that PBP3_PARA_ is the predominant transpeptidase supporting cell division in *K. pneumoniae* at low pH.

### Loss of PBP1b or PBP3_PARA_ differentially impacts Beta-lactam resistance at low pH in *Klebsiella*

The increased expression of genes encoding PBP2_PARA_ and PBP3_PARA_ and other cell wall maintenance proteins at low pH raised the possibility that these proteins might contribute to acid-dependent Beta-lactam resistance. To address this hypothesis, we generated strains individually defective in a subset of *K. pneumoniae* genes implicated in cell wall or shape maintenance and assayed their response to beta-lactam antibiotics during growth at acidic and neutral pH conditions. In addition to genes that displayed pH-dependent expression, we also evaluated the role of class A PBPs in *K. pneumoniae* response to beta-lactams across pH conditions, as these are known to influence antibiotic response in *E. coli.* While most individual mutations did not significantly impact acid-dependent-Beta-lactam resistance, loss of PBP1b or PBP3_PARA_ altered sensitivity to Beta-lactams at low pH (Fig 2A-D, Supp table 6).

Loss of PBP1b (Δ*mrcB*) most dramatically affected acid-dependent resistance to Cephalexin. A Δ*mrcB* mutant displayed only a 4-fold increase in cephalexin MIC at low pH versus at neutral pH, while the WT has a 64-to 128-fold increase in resistance under the same conditions. The Δ*mrcB* mutant also no longer exhibited acid-dependent resistance to the generalist (AMP) and PBP2-targeting-(DOR, MER) beta-lactams, (Fig 2 A-D, Sfig 6, Supp table 6).The increased beta-lactam sensitivity of a PBP1b deletion mutant at low pH is likely due to loss of the enzyme activity under this condition. Deletion of PBP1b’s cognate outer membrane activator lpoB (Δ*lpoB*) results in similar sensitization levels at low pH as a PBP1b mutant (Sfig 6, Supp Table 6). Defects in PBP1b also led to a nearly 10-fold increase in susceptibility to Cefsulodin in both acidic and neutral pH compared to WT (Sup Table 6). This pH-independent increase in sensitivity of the ΔPBP1b mutant is consistent with data suggesting that PBP1a is the primary Class A target of Cefsulodin^30,31^.

Loss of PBP3_PARA_ differentially impacted resistance to beta-lactams. The PBP3 _PARA_ null mutant no longer displayed acid-dependent resistance to Ampicillin and Cefsulodin at low pH (Fig 2A & B). In fact, a PBP3 _PARA_ deletion mutant was more sensitive to Ampicillin at low pH than at neutral pH (Fig 2A). When challenged with PBP3-targeting Beta-lactams (CEX, AZT, PIP), the PBP3_PARA_ deletion mutant still retained mild acid-dependent resistance but was 2-to 8-fold more sensitive at low pH than WT (Fig 4 D, 5 B). Surprisingly, the PBP3_PARA_ null mutant exhibited increased resistance when challenged with PBP2-targeting beta-lactams at low pH. The Meropenem and Doripenem MICs obtained for the PBP3_PARA_ null mutant were 12-16-fold higher at acidic pH versus neutral pH. For comparison, WT cells only exhibited a 2-4-fold increase in MIC to these two drugs at acidic pH (Fig 4C, 5A). Despite the ∼10-fold increases in expression under acidic conditions, loss of *mrdA2,* encoding PBP2, had no significant impact on Beta-lactam susceptibility across pH values.

**Fig 5.**
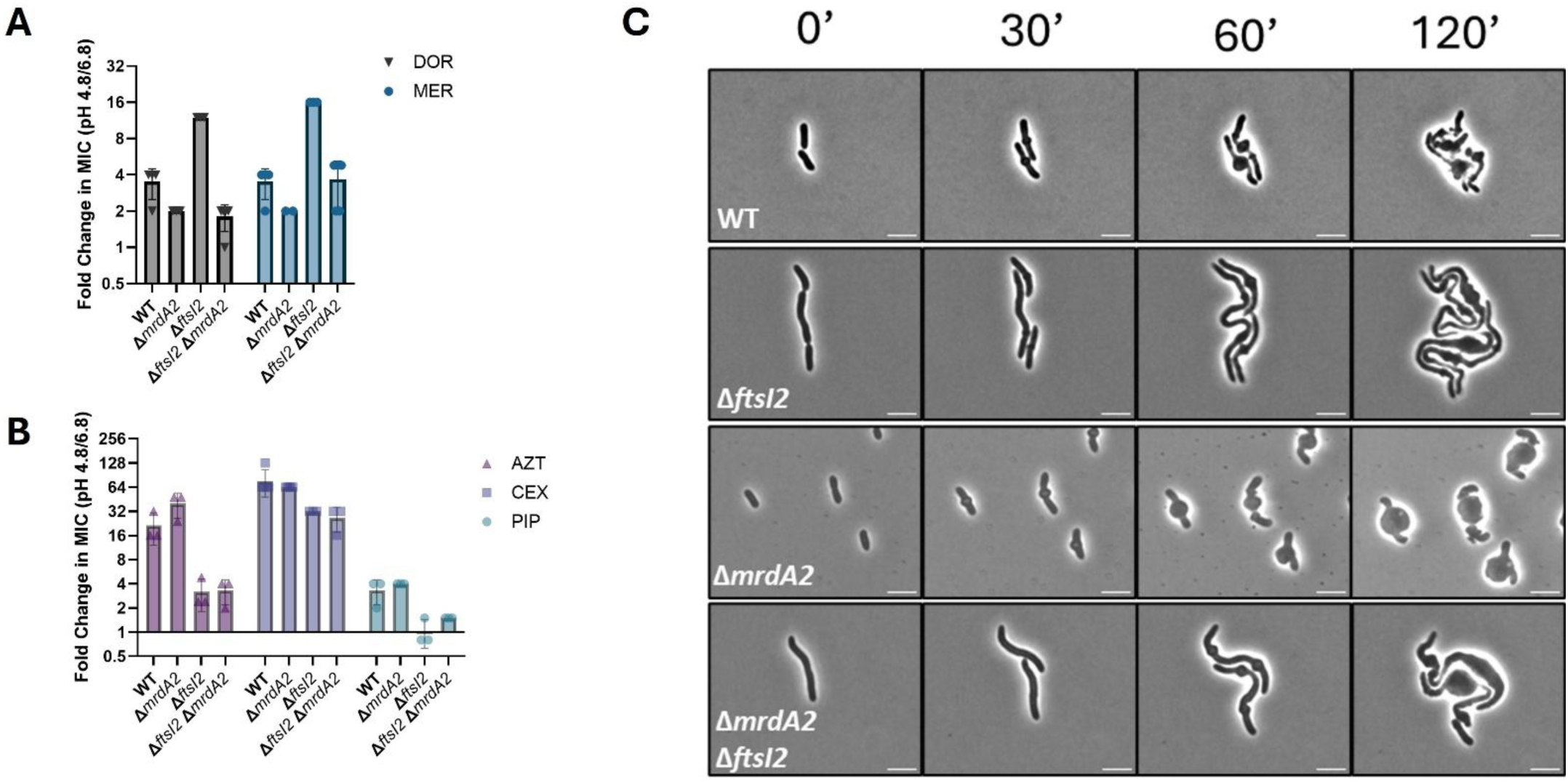
Loss of either PBP1b or PBP3_PARA_ results in differential impacts to Beta-lactam resistance at low pH. (A-D) Fold change in MICs at pH 4.8 relative to pH 6.8 shown for generalist (A), class A-targeting (B),PBP2-targeting (C), and PBP3-targeting Beta-lactams. For comparison, data for MER and CEX are re-graphed from Fig 4. Data are presented as median values with range. (E) Micrographs from time-lapse microscopy of WT, Δ*ftsI2,* Δ*mrdA2,* and Δ*mrdA2* Δ*ftsI2* growing in the presence of MER at low pH. Microscopy pads comprised of LB buffered with MMT to pH 4.8 + 0.2 µg/mL MER + 1.5 µM propidium iodide + 1% agarose. The MER concentration reflects 1X MIC for WT, ∼0.25X MIC for Δ*ftsI2, ∼*2X MIC for Δ*mrdA2*, and ∼0.5X MIC for Δ*mrdA2* Δ*ftsI2*.

### Increased acid-dependent resistance to PBP2-targeting compounds in the PBP3_PARA_ null mutant requires PBP2_PARA_

We reasoned that the unexpected increase to PBP2-inhibitors at low pH observed in the Δ*ftsI2* mutant could be influenced by the presence of PBP2_PARA_. To address this possibility, we generated a double deletion strain lacking both class B paralogs (Δ*ftsI2* Δ*mrdA2*) and assayed its response to beta-lactams with high specificity for PBP2 or PBP3 at both neutral and acidic pH. Deletion of both class B PBP paralogs eliminated the dramatic increase in resistance at low pH (4.8), restoring WT-level responses to both Doripenem and Meropenem. When challenged with PBP3-targeting Beta-lactams, the double mutant class B paralog mutant phenocopied the Δ*ftsI2* mutant, displaying similar reductions in resistance at low pH (Fig 5). Together these data suggest that both PBP2_PARA_ and PBP3_PARA_ are involved in mediating resistance to PBP2-inhibitors, whereas resistance to PBP3-targeting Beta-lactams is primarily facilitated by PBP3_PARA_.

## Discussion

As observed in *E. coli,* beta-lactam resistance in *K. pneumoniae* increases substantially during growth in acidified medium (Fig 1A). Importantly, we observed pH-dependent increases in beta-lactam resistance in LB as well as in a more physiologically relevant medium, artificial urine (Fig 1, Sfig 1). We find that the class A PBP, PBP1b, and the acid-responsive paralogous copy of the class B PBP, PBP3_PARA_, are major genetic contributors to acid-mediated beta-lactam resistance in *K. pneumoniae.* Disruption of *mrcB*, the gene encoding PBP1b, eliminates acid-mediated resistance to the class B PBP inhibitors, Meropenem, Doripenem, and Cephalexin. Loss of PBP3_PARA_ differentially impacts beta-lactam resistance, with mutants showing sensitization towards Ampicillin, Cefsulodin, and all PBP3-targeting beta-lactams but increased resistance towards PBP2-targeting antibiotics.

While both *E. coli* and *K. pneumoniae* demonstrate acid-dependent beta-lactam resistance, factors influencing this phenomenon differ between the two organisms. Acid-mediated beta-lactam resistance in *E. coli* appears to be solely due to intrinsic properties of the PBP enzymes, particularly PBP1b^3^, while in *K. pneumoniae* the contribution of PBP1b is supplemented by pH-dependent induction of genes encoding paralogous class B PBPs. Although we only observed strong changes to antibiotic susceptibility in a PBP3_PARA_ null mutant, loss of PBP2_PARA_ impacted the resistance to PBP2-targeting antibiotics in a PBP3_para_ null background, as well as the morphological response at low pH (Fig 5 C). This result suggests that PBP2_PARA_ plays a role in maintaining PG and cell shape at low pH. Together, our findings support a modular model of cell wall synthesis within the *Enterobacteriaceae* family where apparent function redundancy amongst cell wall synthesis enzymes is protective by enabling PG synthesis across a range of pH conditions.

### The Class A PBP, PBP1b, is important for acid-dependent beta-lactam resistance in *K. pneumoniae*

PBP1b is required for resistance to PBP2- and PBP3-targeting beta-lactams and for growth at low pH ( <5) in *E. coli*^3^. In contrast, deletion of PBP1b in *K. pneumoniae* did not significantly impair growth in acidified culture medium (pH 4.8), suggesting that there are key differences in how PBP1b influences growth in acidic environments between *E. coli* and *K. pneumoniae* (Sfig 5). While expression of *mrcB,* encoding PBP1b, was not influenced by pH, loss of PBP1b impaired *K. pneumoniae’s* resistance to Ampicillin, Doripenem, Meropenem, and Cephalexin at low pH (Fig 2, Supp Table 5). Consistent with MIC data, a PBP1b-deficient strain exhibits faster killing kinetics than a WT strain in the presence of Cephalexin at low pH (Fig 2 E &F). While filamentation is a characteristic morphology associated with division-targeting beta-lactams^32,33^, at low pH a PBP1b-deficient *K. pneumoniae* strain is sensitive to Cephalexin at concentrations that do not elicit filamentation (Fig 4E). The time frame for lysis of the PBP1b mutant is rapid with more than 50% of the population exhibiting lysis or loss of membrane integrity as denoted by propidium iodide staining within 90 minutes of exposure to Cephalexin at low pH (Fig 2F).

Notably, at neutral pH loss of PBP1b or its cognate outer membrane activator, LpoB, mutant did not significantly alter MICs across beta-lactam classes (Sup Table 5), suggesting that PBP1B specifically mediates beta-lactam resistance at low pH in *K. pneumoniae*. This finding differs from what has been observed in *E. coli* where loss of either PBP1b or LpoB increases sensitivity to multiple classes of beta-lactams at neutral pH^34,35^. This result is also consistent with prior data suggesting that PBP1a does not function optimally under acidic conditions and therefore cannot compensate for loss of PBP1b activity at low pH^3^. Our data demonstrating that a PBP1b mutant is more sensitive to PBP2- and PBP3-targeting beta-lactams is consistent with prior studies reporting interactions between PBP1b and key division and elongasome proteins^36,37,29^. Based on these data, we propose that PBP1b supports cell wall integrity at low pH through interactions supporting other cell division proteins.

### Class B PBP paralogs differentially modulate pH-dependent changes in beta-lactam resistance

Although, low pH induced expression of genes encoding both *K. pneumoniae’s* PBP2 and PBP3 paralogs (Table 1), only loss of *ftsI2,* encoding PBP3_PARA_, sensitized cells to beta-lactams during growth in acidified medium. These data are consistent with observations in *S.* Typhimurium that loss of its PBP3 paralog, PBP3_SAL_sensitizes cells to beta-lactams targeting division^38^.Defects in *mrdA2*, encoding PBP2_PARA_, did not significantly impact beta-lactam resistance at either acidic or neutral pH. In *K. pneumoniae* loss of PBP3_PARA_ at low pH sensitized cells to not only PBP3 inhibitors, but also to a Cephalosporin targeting class A PBPs and the penicillin, Ampicillin (Figure 4 A&B).

While loss of PBP3_PARA_ generally sensitized *K. pneumoniae* to beta-lactams under acidic conditions, it increased resistance to compounds targeting PBP2 at pH 4.8 up to 12-to 16-fold (Figure 5A). This increase in resistance in PBP3_PARA_ deficient cells was eliminated in a double mutant lacking both PBP3_PARA_ and PBP2_PARA_ (Δ*ftsI2* Δ*mrdA2*). Under acidic conditions the double class B paralog mutant exhibited WT-level MICs when challenged with Meropenem or Doripenem (Figure 5).

It is unclear why the loss of PBP3_PARA_ renders cells resistant to PBP2-targeting compounds. Morphological data of WT *K. pneumoniae* challenged with Meropenem at low pH reveals hybrid morphologies, with cells displaying signatures of both divisome and elongasome inhibition. Cells first undergo lengthening and then display large, rounded septal bulges (Fig 5C). These results could reflect altered target binding by Meropenem at low pH, where PBP3/PBP3_PARA_ are also inhibited along with PBP2/PBP2_PARA_. Alternatively, physical interactions between class B PBPs within the elongasome and divisome have been reported in *E. coli*^39^, and data suggests that the protein components of the PG synthesis complexes in *Salmonella* vary depending on whether the canonical or paralogous class B PBP is present in the complex^29,40^. Alterations to interactions between class B PBP paralogs and other cell wall synthesis proteins due to beta-lactam binding or in the class B PBP paralog mutant backgrounds could influence morphology and antibiotic resistance. Additional work is needed to fully understand the complex role that the class B PBP paralogs play in resistance to PBP2-targeting beta-lactams.

### PBP3_PARA_ is critical for *K. pneumoniae* division at low pH

Our data suggest that PBP3_PARA_ is the major transpeptidase supporting cross wall synthesis at low pH. Loss of PBP3_PARA_ did not impact cell length at neutral pH but impaired division at low pH, resulting in filamentous cells. However, these acid-cultured PBP3_PARA_-deficient filamentous cells were able to resume division and return to WT-lengths if they were transitioned to neutral pH (Sfig3), suggesting that the canonical PBP3 is unable to fully compensate for the loss of PBP3_PARA_ at low pH. We attempted to disrupt the canonical PBP3 under acidic culture conditions where PBP3_PARA_ is likely the dominate division-specific transpeptidase; however, despite multiple attempts, we were unsuccessful. Moreover, a small percentage of Δ*ftsI2* cells maintained WT-lengths at low pH, suggesting that the canonical PBP3 retains limited, but critical activity at low pH (Sfig 3). These data differ from *Salmonella* where the canonical PBP3 and PBP3 paralog PBP3_SAL_ are individually dispensable at low pH. The canonical PBP3 is capable of supporting growth at both low and neutral pH, but PBP3_SAL_ is unable to support division at neutral pH^28^. Our results support a model in which *K. pneumoniae* PBP3 functions optimally at neutral pH while PBP3_PARA_ functions predominately at acidic pH, with both working together to ensure that division remains robust across pH conditions.

### Expansion and pH specialization of *K. pneumoniae* cell wall synthesis enzymes provide novel therapeutic targets

Identification of paralogous class B PBPs has thus far only been described in pathogens ^27–29,41^, but the role of these paralogous cell wall synthesis proteins in pathogen physiology remains largely uncharacterized. In *Salmonella,* the acid-responsiveness of class B PBP paralogs raised speculation that they could be critical to support bacterial survival during intracellular growth within acidified host immune cells. However, loss of the class B PBP paralogs did not impair replication within macrophages or attenuate virulence in a mouse infection model^28^.

In *Klebsiella pneumoniae,* our data suggests that class B PBP paralogs are part of a general strategy of functional redundancy to preserve critical enzymatic activity across a wide range of environmental conditions. Colonization of a host exposes a bacterial pathogen to a wide range of pH conditions^42–44^. We demonstrate that the acidification of growth medium to physiologically-relevant levels is sufficient to increase resistance to beta-lactams. The class B PBP paralogs modulate that resistance in part, suggesting that these pathogen-specific proteins may provide novel therapeutic targets. We propose that future screens for new antimicrobial targets factor environmental pH as a variable to identify novel compounds that can bind to acid-responsive targets such as the Class B PBP paralogs. Altogether, our results highlight the need to evaluate pathogen physiology in diverse environments beyond the standard laboratory conditions.

## Acknowledgements

We thank members of the Levin lab for their feedback and discussion regarding this study, with a special thanks to Kyra Raines and Dr. Sarah Anderson for critical reading of this manuscript. We are also grateful to David Rosen and his group for the gift of TOP52 and welcome advice on genetic manipulation of *K. pneumoniae.* This work has been funded by the NIH through an R35-GM127331 award to P.A.L.

## Contributions

S.B.: Conceptualization, Investigation, Formal analysis, Methodology, Visualization, Writing-original draft, Writing – review and editing.

P.A.L.: Conceptualization, Funding acquisition, Resources, Supervision, Writing – review and editing.

## Data availability

All raw and processed data required to evaluate the conclusions of this study are included in the text or supplemental data. The RNA sequencing data for gene expression studies at acidic and neutral pH have been deposited in the NCBI GEO repository: GSE291667.

## Methods

### Bacterial strains, growth conditions, and antibiotics

Bacterial strains and plasmids used in this study are listed in Supplementary table 1 and Supplementary table 2. Strains were routinely cultured in Lysogeny Broth (LB – 1% Tryptone, 0.5% Yeast extract, and 1% NaCl) in large glass tubes at 37 °C with 200 RPM aeration. The pH of LB was adjusted with 1:10 MMT buffer (1:2:2 molar ratio of D/L-malic acid, MES, and Tris base). Artificial urine (AU) was prepared as described^18^, and cells were cultured in either base AU or AU supplemented with all twenty amino acids at a final concentration consistent with what is used in EZ rich defined medium (https://www.genome.wisc.edu/resources/protocols/ezmedium.htm). The pH of AU or AU+AA was adjusted with MMT buffer immediately prior to use. For ease and clarity, media and amino acid composition is listed in Supp table 8.

Where indicated cultures were supplemented with Kanamycin (50 µg/mL), Spectinomycin (100 µg/mL), Chloramphenicol (50 µg/mL), or IPTG (0.1 mM). The pRDC3 plasmid (p15A ori, SpecR, Plac) was a gift from Fabrizio Arigoni. Plasmid pSDB8 was generated from pRDC3 using the IVA cloning method to insert *ftsI2* under control of the Plac promoter^45^. Primers were acquired from Integrated DNA Technologies (Coralville, IA) and sequences used are listed in Supp Table 3. Chemically competent *E. coli* TOP10 (Invitrogen) were used to assemble vectors via IVA. Plasmid sequence was confirmed via whole plasmid sequencing at Plasmidsaurus prior to transformation into *K. pneumoniae* strains.

For growth studies, cultures were started in LB supplemented with MMT buffer to the indicated pH and grown in large glass tubes at 37 °C with 200 RPM aeration. Cultures were harvested once they reached mid-exponential phase (OD_600_ =0.2-0.6), concentrated to an OD_600_ =1.0, and serially diluted 10-fold. 1.5 µL of the 10^-5^ dilution was used to inoculate wells of sterile 96-well plates containing 150 µL of LB + MMT at indicated pH. The plate was covered with a Breathe-Easy strip (Diversified Biosciences) and growth was monitored in a BioTek Synergy H1 plate reader with the following settings: 37 °C, continuous double orbital shaking at 282 cpm, with OD_600_ readings taken every 10 minutes.

For *ftsI2* cell size complementation studies, un-induced overnight cultures of strains carrying either pRDC3 (Empty vector, EV) or pSDB8 were back-diluted 1:500 into 3 mL of fresh LB buffered to either pH 4.8 or pH 6.8 containing Spectinomycin (100 µg/mL). Cultures were grown at 37 °C with 200 RPM aeration for 1.5 hrs. Expression of *ftsI2* was induced with the addition of 0.1 mM IPTG and cultures were allowed to continue growing until early exponential phase (OD_600_ = 0.1-0.3) was reached (∼40-90 additional minutes). Cultures were back-diluted into fresh buffered media containing Spectinomycin (100 µg/mL) and 0.1 mM IPTG to an OD_600_ ∼= 0.005. Growth was monitored every 30 minutes, and cells were harvested when cultures reached early exponential phase (OD_600_ = 0.1-0.3). Cells were spotted onto media pads (LB + MMT (pH 4.8 or 6.8) + Spectinomycin (100 µg/mL) + 0.1 mM IPTG + 1% agarose) and imaged live with phase microscopy.

The following antibiotics were used for antibiotic susceptibility screenings. Ampicillin sodium salt, Aztreonam, Cephalexin hydrate, Cefsulodin sodium salt hydrate, Meropenem Trihydrate, and Piperacillin sodium salt were obtained from Sigma Adrich (St. Louis, MO). Carbenicillin sodium salt was from EMD Millipore (now Millipore Sigma, Germany). Doripenem was obtained from Cayman Chemicals (Ann Arbor, MI).

### Genetic manipulation

Gene disruptions were carried out in the *K. pneumoniae* TOP52 background using a Lambda Red vector modified to carry a Spectinomycin resistance cassette as described previously ^46,47^. Briefly, DreamTaq Polymerase (Thermo Scientific) was used to PCR amplify the Kanamycin resistance cassette from the template plasmid pKD4 with primers containing 40-60 bp of homology to the chromosomal targets. Primers were designed to leave the start and stop codons of the targeted gene intact. In cases where the two gene sequences overlapped, primers were designed to avoid disruption to any downstream genes. Linear PCR products were purified using a Purelink PCR purification kit (Invitrogen), and 2,000-3,000 ng of purified linear fragments were electroporated into *K. pneumoniae* cells grown in the presence of 0.7% L-arabinose to induce expression of the Lambda Red recombinase system from the pKD46S plasmid. Electroporated cells were recovered in SOC (Lloyd’s recipe – 2% Tryptone, 0.5% yeast extract, 10 mM NaCl, 2.5 mM KCl, 10 mM MgSO_4_, 20 mM glucose) for 4-8hrs at 30°C. Deletion mutants were selected for by plating transformation culture onto LB agar plates containing Kanamycin (50 µg/mL) and incubated at 37°C. Mutants were confirmed by colony PCR and sequencing as necessary. For marker-less deletions, cells were transformed with pCP20 at 30°C in LB + Chloramphenicol (50 µg/mL) to express the FLP to excise the FRT-flanked Kanamycin cassette. After excision, cells were cultured at 42°C to promote loss of pCP20. Kanamycin and Chloramphenicol sensitivity was confirmed for strains prior to use.

### RNA sequencing

*K. pneumoniae* TOP52 or *E. coli* MG1655 was cultured overnight in LB buffered to either pH 4.8 or 6.8 with MMT buffer. Overnight cultures were back diluted 1:100 into fresh media and cultured at 37°C with 200 RPM aeration until cultures reached early exponential phase (OD_600_ =∼0.2). ∼ 9.6 x 10^8^ cells were transferred to 15 mL conical Falcon tubes and pelleted by centrifugation at 4°C (4,000 RPM for 10 mins). The supernatant was discarded, cell pellets were resuspended in 350 uL of RNAwiz, and the RNA was extracted following the protocol for the Ribopure-Bacteria RNA extraction kit [Invitrogen]. RNA was eluted in 50 uL of elution solution. Purified RNA was subjected to a DNase I treatment to remove any contaminating chromosomal DNA. RNA sequencing and differential gene expression analysis was performed by MiGS (now SeqCenter). Quality control and adapter trimming of reads was performed with bcl2fastq. Read mapping was performed with HISAT2, and read quantification was performed using Subread’s featureCounts. Read counts were normalized in R using edgeR’s Trimmed Mean of M values (TMM) algorithm, and subsequent values were converted to counts per million (CPM). Differential expression analysis was performed using edgeR’s Quasi-Linear F-Test (qlfTest) functionality against treatment groups, and DEGs were considered as those with |logFC| > 1 and p < .05. Three biological replicates were submitted per pH condition.

### Antibiotic susceptibility testing

Cells were cultured from a single colony in LB at the indicated pH to exponential phase (OD_600_ =∼0.2-0.6) at 37 °C with 200 RPM aeration. For MICs in AU or AU+AA, overnight cultures were back-diluted 1:100 into fresh media and cultured at 37 °C with 200 RPM aeration at the indicated pH to exponential phase (OD_600_ =∼0.2-0.6). Harvested cells were concentrated to an OD_600_ =1 and serially diluted 10-fold. 1.5 µL of the 10^-2^ dilution was used to seed wells containing 150 µL of LB broth + MMT (matching pH) in sterile 96-well microtiter plates containing a range of two-fold dilutions of beta-lactam antibiotics. Plates were sealed with a Breathe-Easy strip (Diversified Biotech) and incubated for 20 hours at 37°C with 200 RPM aeration. The minimal inhibitory concentration (MIC) was determined by visual inspection and reflects the lowest drug concentration that completely inhibited bacterial growth.

### Terminal morphology Assessment

At the end of 20 hours for antibiotic susceptibility tests, cells were taken from wells following growth at either neutral or acidic pH with the indicated antibiotic concentrations. 5 µL of cells were spotted onto LB + 1% agarose and imaged by phase contrast microscopy.

### Microscopy, cell size analysis, and time-lapse Imaging

All phase contrast and fluorescence images were acquired on either a Nikon Ti-E inverted microscope or a Ti2-E inverted microscope (Nikon Instruments). The Ti-E is equipped with a 100X Plan Apo oil objective (N.A. = 1.45 Ph3 objective, X-Cite 120 LED light source (Lumen Dynamics) and an OrcaERG CCD camera (Hammamatsu Photonics, Bridgewater, N.J.) Filter cubes were purchased from Chroma Technology Corporation. The Ti2-E inverted microscope is equipped with a 100X Plan Apo oil objective (N.A. = 1.45, Ph3 objective, Lumencor Sola LED light engine (Lumencore), and an orca-Fusion sCMOS camera (Hammamatsu Photonics). Filter cubes were purchased from Chroma Technology Corporation. Both microscopes have an enclosed temperature-controlled enclosure to pre-heat the objective to 37 °C (OXO enclosure) prior to experiments. Nikon Elements software was used for image capture, and analysis was performed using the MicrobeJ plug-in in ImageJ.

For cell size analysis, cells were grown from a single colony in LB with MMT buffer at indicated pH at 37 °C and 200 RPM aeration to exponential phase (OD_600_ = 0.2-0.4) and back-diluted to an OD_600_ = 0.005 into 7 mL of pre-warmed LB + MTT (same pH). Growth was monitored via optical density, and when cultures reached early exponential phase (OD_600_ =0.1-0.2) cells were harvested. 5 µL of cells were spotted onto LB + MTT (same pH)+ 1% agarose pads and imaged by phase contrast microscopy. Cell morphology and size dimensions were determined using the MicrobeJ^48^ plug-in for ImageJ.

For time-lapse imaging, cultures were grown from a single colony in LB with MMT buffer at indicated pH at 37 °C and 200 RPM aeration to exponential phase (OD_600_ = 0.1-0.4). 5 µL of cells were spotted onto LB + MTT (same pH)+ 1% agarose pads that contain antibiotics at indicated concentrations and 1.5 µM propidium iodide (PI) for monitoring loss of membrane integrity. Phase images were obtained every 2 minutes, and fluorescent images were collected every 10 minutes (Chroma AT560/40X/ Chroma AT600 DC/ Chroma at635/60nm with 100 mS exposure/10% intensity). For each experiment a total of 2-5 different XY coordinates were monitored throughout the duration of the experiment. Quantification of PI^+^ cells was done using the MicrobeJ^48^ plug-in for ImageJ. Briefly the number of cells in the phase image and the number of corresponding fluorescent cells were quantified. Data was reported as either percentage of PI^+^ cells (number of PI^+^ cells/total number of cells *100) or percent alive (number of cells-PI^+^ cells/total number of cells*100).

## Notes

https://www.ncbi.nlm.nih.gov/geo/query/acc.cgi?acc=GSE291667

### Competing Interest Statement

The authors have declared no competing interest.

### Summary of Updates

This version has been updated to include new experimental data included in Sfig1 and 6 and in Supplemental tables 5-8.

