## Supplemental tables 1-3, Supplemental figures 1-6 for "pH-dependent beta-lactam resistance in *Klebsiella pneumoniae* is mediated by paralogous class B PBPs and the class A PBP, PBP1b"

**Supplementary Table 1.** Strains used in this study.

| Strain ID | Genotype | Source |
|---|---|---|
| PAL4422 | *Klebsiella pneumoniae* TOP52 | [1,2] |
| PAL4649 | NTUH-K2044 | [3] |
| PAL4615 | ATCC 43816 | [2] |
| SDB583 | *K. pneumoniae* TOP52 Δ*bla*::kan | This study |
| SDB595 | *K. pneumoniae* TOP52 Δ*mrcA*::kan | This study |
| SDB597 | *K. pneumoniae* TOP52 Δ*mrcB*::kan | This study |
| SDB624 | *K. pneumoniae* TOP52 Δ*pbpC*::kan | This study |
| SDB676 | *K. pneumoniae* TOP52 Δ*mrdA2*::kan | This study |
| SDB689 | *K. pneumoniae* TOP52 Δ*ftsI2*::kan | This study |
| SDB588 | *K. pneumoniae* TOP52 Δ*dacA*::kan | This study |
| SDB591 | *K. pneumoniae* TOP52 Δ*dacD*::kan | This study |
| SDB700 | *K. pneumoniae* TOP52 Δ*ftsI2*::frt Δ*mrdA2*::kan | This study |
| SDB757 | *K. pneumoniae* TOP52 Δ*lpoB*::kan | This study |
| PAL2036 | *Escherichia coli* MG1655 | [4] |

**Supplementary Table 2.** Plasmids used in this study.

| **Plasmid ID** | **Genotype/Annotation** | **Source** |
|---|---|---|
| pRDC3 | Empty vector, Plac, SpecR | Gift from Fabrizio Arigoni |
| pSDB8 | pRDC3, *Plac::ftsI2* | This study |
| pKD46S | Lambda Red Recombinase, SpecR | [5,6] |
| pKD4 | Source of Kanamycin Resistance cassette | [6] |
| pCP20 | Source of FLP | [6] |

**Supplementary Table 3.** Primers used in this study.

| Primer pair | Usage | Fwd primer seq (5' - 3') | Rvs primer seq (5' - 3') |
|---|---|---|---|
| PP_28 | generate *blaSHV-1* KO fragment | gcctttatcggccctcactcaaggatgtattgtggttatggtgtaggctggagctgcttc | tgctacgagccggataacgcgcgcggccaccgccgggttacatatgaatatcctccttag |
| PP_27 | Confirm size of blaSHV-1 locus | atttgtcgcttctttactcg | gtgctacgagccggataac |
| PP_51F/PP_30R | generate *mrdA2* KO fragment | cgttggcttttccagtgattcaggtgatgtaccgctaatggtgtaggctggagctgcttc | actatgaacgatgcgcctgcggaataagtcctgtcgcttacatatgaatatcctccttag |
| PP_29 | confirm size of *mrdA2 locus* | acgatccctctaacatgctg | cgatgggacaaataaccaattac |
| PP_59 | generate *mrcA* KO fragment | ataaactgcccaaataaaactaaatgggaaaacaccggtggtgtaggctggagctgcttc | agcagcaataaaaaagcgcccgcaggcgctttttttgtcacatatgaatatcctccttag |
| PP_58 | Confirm size of *mrcA* locus | ggctatatcaatgcacctgg | cgtccacgctaagcagca |
| PP_70 | generate *mrcB* KO fragment | atcaggcctttgcgcctgaacgttgcggagaaaaagcatggtgtaggctggagctgcttc | tactttcgtattaaggggtctggtcgtgcggtcaaaatcacatatgaatatcctccttag |
| PP_56 | confirm size of *mrcB* locus | ttgagagatttcttcttcctctg | cggtcgaaggctgaatt |

| | | | |
|---|---|---|---|
| PP_82_1 | generate *pbpC* KO fragment | aggagcggccagcggtccgctcaccgtgacgccgtaaatggtgtaggctggagctgcttc | ttgaaaacacaaaccgccaacaacaaatttaccctggttatatgaatatcctccttag |
| PP_82 | confirm size of *pbpC* locus | agcatatcgagttccgcg | cgattgtatggcctggcg |
| PP_105 | generate *ftsI2* KO fragment | ctgttaaatcagttggttgcaaaacaatgccggagctatggtgtaggctggagctgcttc | gcgaaagtgtagcgctattcgcccctgcagaccactgttacatatgaatatcctccttag |
| PP_106 | confirm size of *ftsI2* locus | gttaagtggttcaacgagtc | ggtgttgctgaaagggc |
| PP_64 | generate *dacA* KO fragment | ctttttaactccatcacggatgccgttgttctgaccatggtgtaggctggagctgcttc | tatggggatggaaatcacactttcaagtgttcgatttttacatatgaatatcctccttag |
| PP_65 | confirm size of *dacA* locus | aatgtcggatgctggcc | gggcgggagttatcaggttac |
| PP_60 | generate *dacD* KO fragment | aaaggacggcggatctgtaaatctagaggatatgccgttggtgtaggctggagctgcttc | cgaaaaaaaaccgcctgcaaagaggcggttggcgtggttacatatgaatatcctccttag |
| PP_61 | confirm size of *dacD* locus | ccctgatatgaaacgcgaag | gagtcgccaccacactg |
| PP_115 | Used to create pSDB8 via IVA recombineering - backbone linearization | agccaagcttgcatgcctgc | gtaatcatggtcatagctg |
| PP_114 | Used to create pSDB8 via IVA recombineering - Amplification of ftsI2 with backbone homology regions added | tatgaccatgattacatggtgctaaagaaa | gcatgcaagcttggcttacagcgccggccc |
| PP_111 | Used to confirm presence of insert into pRDC3backbone | ttgtgtggaattgtgagcgg | Catcagagcagattgtactgag |
| PP_119 | Generate *lpoB* KO fragment | cacgtcgatccatcacaagaaaatatcgttgtacagaaataaggattattgcgacggatggtgtaggctggagctgcttc | cagcacctttacctgaccagattatctcaccggtttgcaccatatgaatatcctccttag |


**References:** 1.Ko, D. C. *et al*. Whole-Genome Sequencing of Klebsiella pneumoniae Isolates to Track Strain Progression in a Single Patient With Recurrent Urinary Tract Infection. *Frontiers in Cellular and Infection Microbiology | www.frontiersin.org* **9**, 14 (2019).

**Supplemental table legends:**

**Supp table 4.** Beta-lactam MICS for *K. pneumoniae* TOP52 in LB across five pH values. Corresponds to heatmap data presented in Fig 1.

**Supp table 5.** Beta-lactam MICs for *K. pneumoniae* TOP52 in artificial urine (AU) and artificial urine supplemented with amino acids (AU+AA) at pH 4.8 and 6.8.

**Supp table 6.** Beta-lactam MICs for *K. pneumoniae* TOP52 and mutants in LB at pH 4.8 and 6.8.

**Supp table 7.** List of differentially regulated genes (DEG) during growth in LB at pH 4.8 versus pH 6.8 in both *E. coli* MG1655 and *K. pneumoniae*.

**Supp table 8.** Composition of Artificial urine and amino acid stocks used to make base AU and AU+AA media.

**Supplemental figure legends**

**Sfig1. *K. pneumoniae* exhibits acid-dependent beta-lactam resistance in a variety of nutrient environments.** Heatmap displaying the median fold change in MIC for the indicated beta-lactams at pH 4.8 compared to pH 6.8 in indicated media conditions (LB = Lysogeny Broth, AU + AA = Artificial Urine supplemented with amino acids, and AU = Artificial Urine). Data obtained in LB is re-graphed from figures 4 and 5 for ease of comparison. Higher values (magenta) indicated increased MIC at low pH, while values of 1 indicate the MIC is unchanged at low pH compared to neutral pH. AMP = Ampicillin, DOR = Doripenem, MER = Meropenem, and CEX = Cephalexin. The exact fold change for CEX could not be determined in AU ± AA as there was growth in all concentrations tested – see also Supplemental table 5.

**Sfig 2. Sequence and domain conservation between canonical and paralogous Class B PBP proteins.** (A & B) Protein sequence conservation between canonical and paralogous copies of PBP2 (A) and PBP3 (B). Purple bars above residues indicate those that are critical for catalysis.

**Sfig 3. Distribution of cell lengths in Δ*ftsI2* populations during growth at low pH.** (A) Histograms showing the distributions of *K. pneumoniae* Δ*ftsI2* cells grown at pH 4.8. A minimum of 50 cells were analyzed for each biological replicate. Bin lengths indicate the maximum length (i.e. a bin length of 5 includes all cells with a length of 1-5 μM). (B) Micrographs of *K. pneumoniae* Δ*ftsI2* cells grown at pH 4.8. Agarose pads are comprised of LB adjusted to pH 4.8 with MMT buffer + 1% agarose. Scale bar = 10 μM.

**Sfig 4. Acid-grown Δ*ftsI2* filamentous cells resume division upon transition from acidic to neutral pH.** *K. pneumoniae* Δ*ftsI2* cells were cultured in LB+MMT, pH 4.8 until early exponential phase ($OD_{600}$ = 0.1-0.2) and then 5 μL were spotted onto either LB + 1% agarose pads buffered to either acidic or neutral pH and containing 1.5 μM propidium iodide to monitor membrane integrity. Scale bar = 10 μM. Representative images of n=2 biological replicates.

**Sfig 5. Loss of *mrcB* does not significantly alter growth at low pH in *K. pneumoniae.*** Growth of *K. pneumoniae* WT (circles) or Δ*mrcB* (open squares) at pH 4.8 (A), pH 5.5 (B), or pH 6.8 (C). Line represents mean values of n=4 biological replicates with error bars denoting standard deviation.

**Sfig 6. Loss of PBP1b or its outer membrane activator LpoB impairs acid-dependent Beta-lactam resistance.** (A) Fold change (FC) in Doripenem (DOR) MICs at pH 4.8 relative to pH 6.8 shown for class A PBP mutants with loss of PBP1a (Δ*mrcA*) or PBP1b (Δ*mrcB*). Data are graphed as median values with range. (B) Fold change (FC) in MIC of indicated beta-lactams at pH 4.8 compared to pH 6.8 for a Δ*lpoB* mutant. AMP = Ampicillin, CEF = Cefsulodin, MER = Meropenem, DOR = Doripenem, and CEX = Cephalexin.

**Supplemental movies 1 and 2.** Corresponds to micrographs in the top panel of Sfig 4. *K. pneumoniae* Δ*ftsI2* cells were cultured in LB+MMT, pH 4.8 until early exponential phase ($OD_{600}$ = 0.1-0.2) and then 5 μL were spotted onto LB + 1% agarose pads buffered to pH 4.8 containing 1.5 μM propidium iodide to monitor membrane integrity. Scale bar = 10 μM. Movie 1 corresponds to the phase contrast images taken every 2 minutes, and movie 2 is the corresponding fluorescent images taken every 10 minutes.

**Supplemental movies 3 & 4.** Corresponds to micrographs in bottom panel of Sfig 4. *K. pneumoniae* Δ*ftsI2* cells were cultured in LB+MMT, pH 4.8 until early exponential phase ($OD_{600}$ = 0.1-0.2) and then 5 μL were spotted onto LB + 1% agarose pads buffered to pH 6.8 containing 1.5 μM propidium iodide to monitor membrane integrity. Scale bar = 10 μM. Movie 1 corresponds to the phase contrast images taken every 2 minutes, and movie 2 is the corresponding fluorescent images taken every 10 minutes.

**Supplemental figures**

Sfig 1

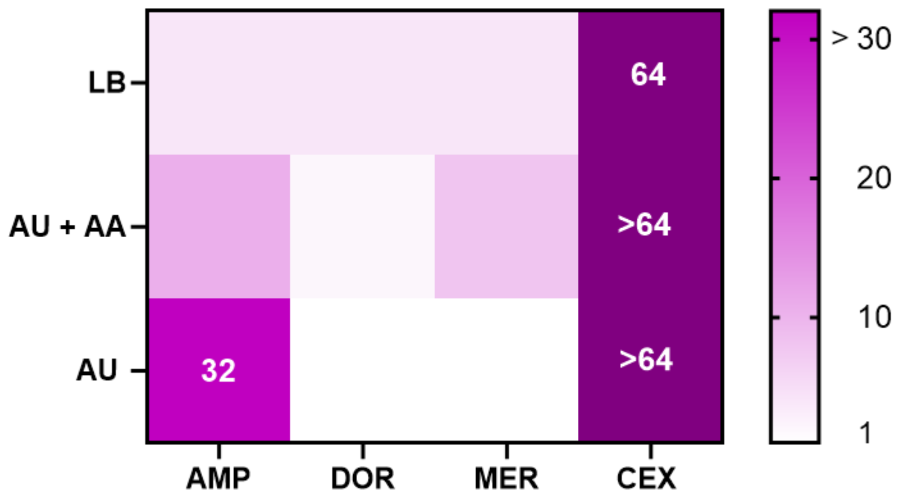

Sfig 2

**A**

```
PBP2       3    LQNSFRDYTAESSLFVRRALVAFLGILLLTGVLIANLYNVQIVRFDDYQTRSNENRIKLV  62
                L++  RD++AE  LF+RRA +AFL +++  GVLI NLY++Q+ + D YQTRSN+N IK++
PBP2_PARA  4    LRDEIRDHSAEEMLFIRRAAIAFLLVVVCFGVLIVNLYHLQVEQHDFYQTRSNQNDIKML  63

PBP2       63   PIPPSRGIIYDRNGTPLALNRTIYQLEMMPEKVDNVQQTLDALRDVVDLTDDDIAGFKKE  122
                PI PSRG+I+DRNG PL  N T+Y+L+++P K+ ++   L  L  +VDLT DDIA F+ +
PBP2_PARA  64   PIAPSRGLIFDRNGIPLVQNITLYRLQVIPSKIPDMAALLQQLTPIVDLTPDDIASFRDD  123

PBP2       123  RARSHRFTSIPVKVNLSEVQVARFAVNQYRFPGVEVKGYKRRYYPYNSALTHVIGYVSKI  182
                   + R+ ++ +K +LS+V+VARFAVN++RFPGV V+ Y++R YPY + L HV+GYVSKI
PBP2_PARA  124  MHHTSRYKAVTLKSDLSDVEVARFAVNEFRFPGVTVESYQQREYPYGAELAHVVGYVSKI  183

PBP2       183  NDKDVDRLDKEGKLANYASTHDIGKLGIERYYEDVLHGQTGYEEVEVNNRGRVIRQLKEV  242
                ND D+ RL K G+  NYA+  +IGK GIE YYE  LHG TGY+EVEV+N GRV+R LKEV
PBP2_PARA  184  NDSDLQRLAKNGEEENYAADRNIGKQGIEGYYEKALHGTTGYQEVEVDNHGRVVRLLKEV  243

PBP2       243  PPQAGRDIYLTLDLKLQQYIETLLAGSRAAVVVTDPRTGAILALVSTPSYDPNLFVDGIS  302
                PP AG+++YLTLDL LQQYIE++L G RAAVVV DPR G +LA+VS+PSYDPN FV GI
PBP2_PARA  244  PPVAGKNLYLTLDLHLQQYIESVLKGQRAAVVVVDPRDGGVLAMVSSPSYDPNPFVKGIG  303

PBP2       303  SKDYSALLNDPNTPLVNRATQGVYPPASTVKPYVAVSALSAGVITRNTSLFDPGWWQLPG  362
                 + Y +LL++P+ PL+NR TQG+YPPASTVKPY+A+SALSAGVIT NT+ F    W LPG
PBP2_PARA  304  YQAYKSLLDNPDRPLINRVTQGLYPPASTVKPYMALSALSAGVITPNTTFFGAPTWTLPG  363

PBP2       363  SEKRYRDWKKWGHGHLNVTKALEESADTYFYQVAYDMGIDRLSEWMSKFGYGHYTGIDLS  422
                +++RYRDW K GHG LNVTKA+EESADT+FYQVA++MGIDR+ EW+SKFGYG  TGIDL+
PBP2_PARA  364  TQRRYRDWLKTGHGMLNVTKAIEESADTFFYQVAFEMGIDRIHEWLSKFGYGQSTGIDLN  423

PBP2       423  EERSGNMPTREWKLKRFKKPWYQGDTIPVGIGQGYWTATPIQMNKALMILINDGVVKVPH  482
                EE +G +P+REWK +  KKPWYQGDTI VGIGQGYW ATPIQM KAL  L+N+G V+ PH
PBP2_PARA  424  EEYAGVLPSREWKQRVHKKPWYQGDTISVGIGQGYWIATPIQMVKALTTLLNNGKVQDPH  483

PBP2       483  LLQSTVEDGKPVPWVQPHE-PPVGDIHSGYWEIAKDGMYGVANRGNGTAHKYFASAPYKI  541
                LL S  +      + QP   P VGD  S YW I ++GMYG+AN+ NGT +K F +APY+I
PBP2_PARA  484  LLYSMKQGNHVERYQQPANLPQVGDPKSPYWGIVRNGMYGMANQPNGTGYKLFHTAPYQI  543

PBP2       542  AAKSGTAQVFGLKANETYNAHRISERLRDHKLMTAFAPYNNPQVAVAMILENGGA-GPAV  600
                AAKSGT+QVF LK N+TYNA  I  RLRDH   T FAPY +P+VA+A+ILENGG  G
PBP2_PARA  544  AAKSGTSQVFSLKQNQTYNAKMIPVRLRDHIFYTLFAPYQHPKVAMALILENGGGDGVVA  603

PBP2       601  GTIMRQILDHIMLGDNNTTLPSENPAVTAGEDQ  633
                G   R ILDHI +    ++  ++ P   + + Q
PBP2_PARA  604  GPTARAILDHIFVPQQASSAAADVPQRDSADAQ  636
```

**B**

```
FtsI       11   KRQEEQANFISWRFALLCGCILLALAFLLGRVAWLQVISPDMLVRQGDMRSLRVQEVSTA  70
                K+ +  A+F   RF LLC  IL  L  LL RV WLQ+ISPD LV+Q DMRSLR + V+
FtsI_PARA  5    KKTKSAASFTPIRFGLLCVAILGCLGLLLVRVGWLQIISPDNLVKQEDMRSLREEPVAVE  64

FtsI       71   RGMITDRSGRPLAVSVPVKAIWADPKELHDAGGVTLDTRWKALADALNMPLDQLATRINT  130
                RGMI+DR GRPLAVSVPV AIW DP+    + GGV    RW+A+A+AL++ L +LA R+ +
FtsI_PARA  65   RGMISDREGRPLAVSVPVSAIWIDPQTTMEKGGVGYGPRWQAMAEALHLNLGELAQRVQS  124

FtsI       131  NPRMRFIYLARQVNPDMADYIKKLKLPGIHLREESRRYYPSGEVTAHLIGFTNVDSQGIE  190
                +P  RF+YLARQ+NP+ A++I KL LPG++LR+ESRR+YP+G V A+L+GFTNVD+QGIE
FtsI_PARA  125  HPHARFLYLARQINPEQAEWIDKLHLPGVYLRDESRRFYPAGHVAANLLGFTNVDNQGIE  184

FtsI       191  GVEKSFDKWLTGQPGERIVRKDRYGRVIEDISSTDSQAAHNLALSIDERLQALVYRELNN  250
                GVEKSF+  LTG+PG R+VRKD++G VIE+I+       AHNL LSIDERLQ +    L+N
FtsI_PARA  185  GVEKSFNAQLTGKPGRRLVRKDKHGNVIENITEVPPVPAHNLQLSIDERLQTVTEDALDN  244

FtsI       251  AVAFNKAESGSAVLVDVNTGEVLAMANSPSYNPNNFAGTAKDTMRNRAITDVFEPGSTVK  310
                AV +NKAESG+AVL+ ++TGE+LAMA+ P +NPNN       D  RNRAI+D FEPGSTVK
FtsI_PARA  245  AVRWNKAESGAAVLIKIDTGEILAMASYPDFNPNNRDSATLDDFRNRAISDTFEPGSTVK  304

FtsI       311  PMVVMTALQRGIVNENTVLNTVPYRINGHEIKDVARYSELTLTGVLQKSSNVGVSKLALA  370
                P+V+MTALQ+GIV  ++V++T P+ ++GH I+DV  Y EL+LTG+LQKSS+ GVS L+LA
FtsI_PARA  305  PLVIMTALQQGIVQPDSVVDTHPFVLDGHRIRDVGYYPELSLTGILQKSSDTGVSHLSLA  364

FtsI       371  MPSSALVDTYSRFGLGKATNLGLVGERSGLYPQKQRWSDIERATFSFGYGLMVTPLQLAR  430
                MP   L+DTY  FG G+ T LGL GE +GL P  + W  ++RATF+FGYGLMVTPLQLA
FtsI_PARA  365  MPVQHLIDTYKAFGFGEPTGLGLTGESAGLMPHHRYWGQLDRATFAFGYGLMVTPLQLAH  424

FtsI       431  VYATIGSYGIYRPLSITKVDPPVPGERVFPESLVRTVVHMMESVALPGGGGVKAAIKGYR  490
                VYATIG +GI RPLSIT++DPPV G RV PES+V +V HMMESVALPGGGG KAA++ YR
FtsI_PARA  425  VYATIGGFGIARPLSITRIDPPVMGTRVMPESIVHSVEHMMESVALPGGGGTKAAVRDYR  484

FtsI       491  IAIKTGTAKKVGPDGRYINKYIAYTAGVAPASHPRFALVVVINDPQAGKYYGGAVSAPVF  550
                +A+KTGTAKK+GPDG+YI+KY+AYTAGVAPAS P+FALVVV+NDP  G YYGGAVSAPVF
FtsI_PARA  485  VAVKTGTAKKIGPDGKYIDKYVAYTAGVAPASRPQFALVVVMNDPSNGSYYGGAVSAPVF  544

FtsI       551  GAIMGGVLRTMNIEPDALATGEKSEFVINQGEGTG  585
                  IMG VLR  N+ PD +  G ++  V++     G
FtsI_PARA  545  SQIMGDVLRLENVMPDGMPQGAENLIVMHDSHPQG  579
```

Sfig 3

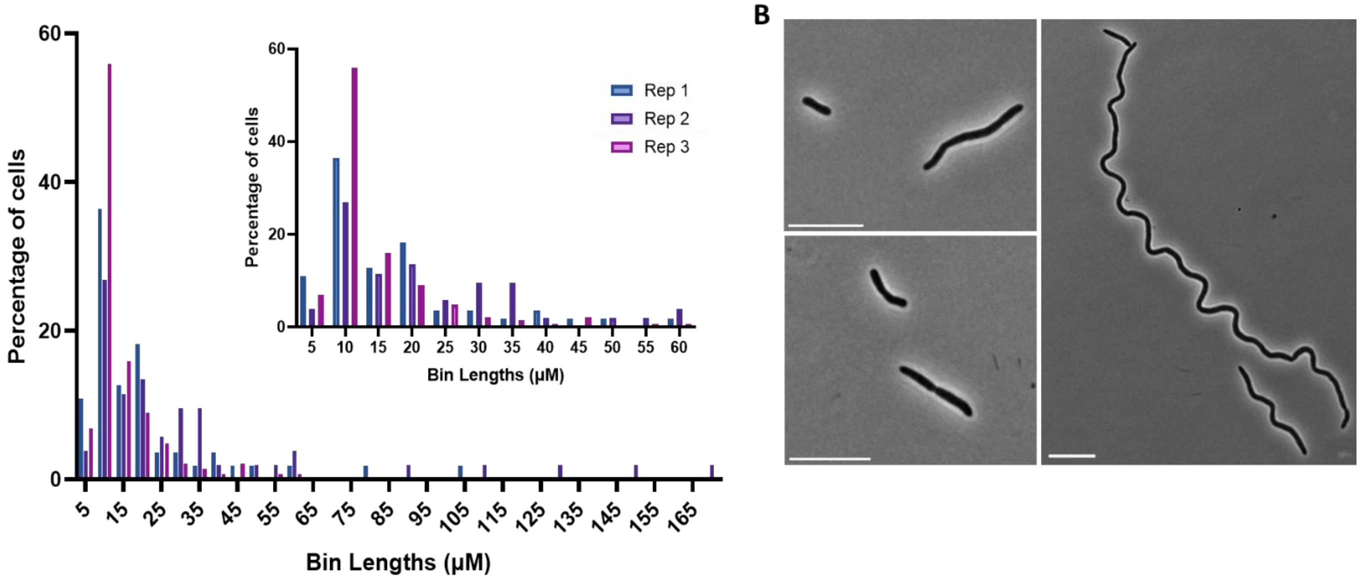

Sfig 4

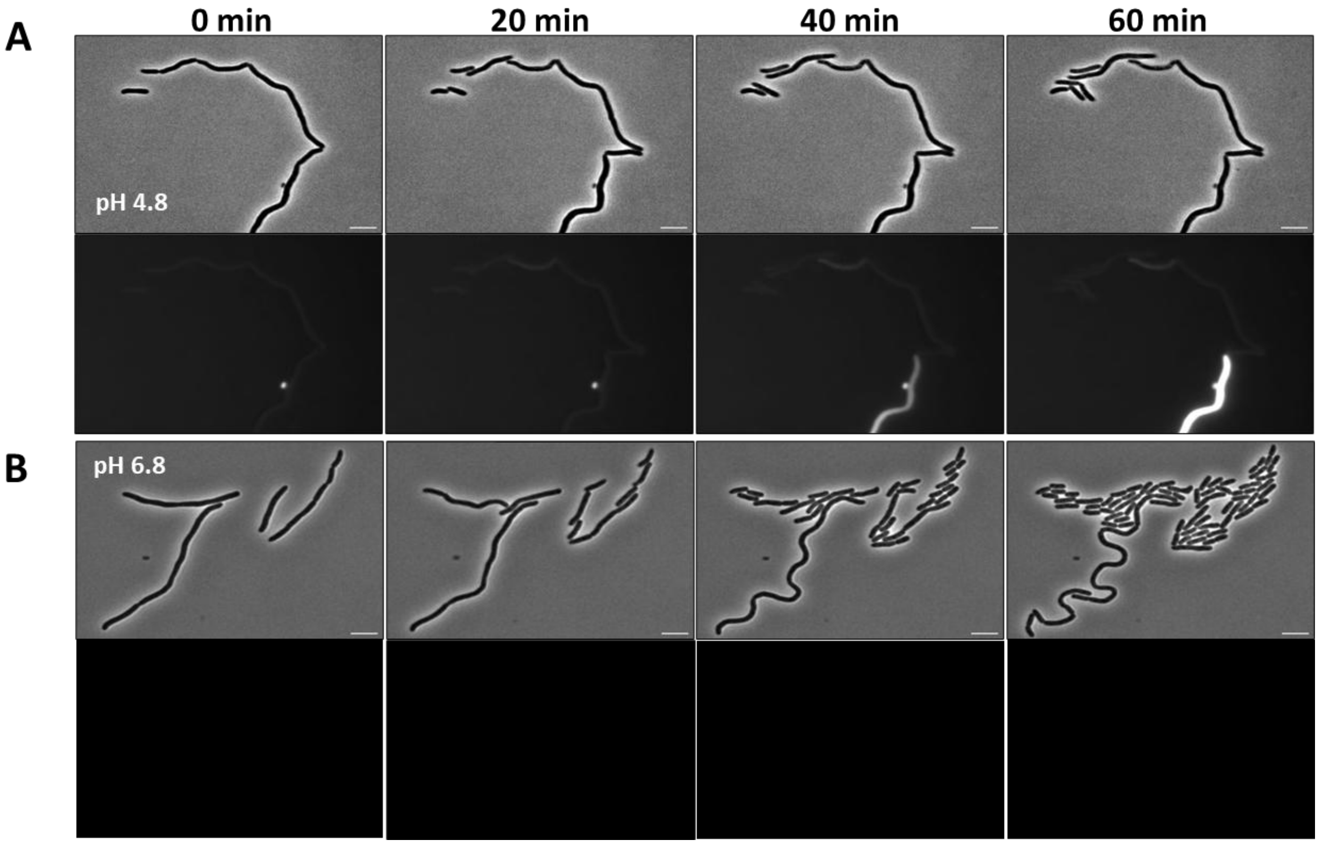

Sfig 5

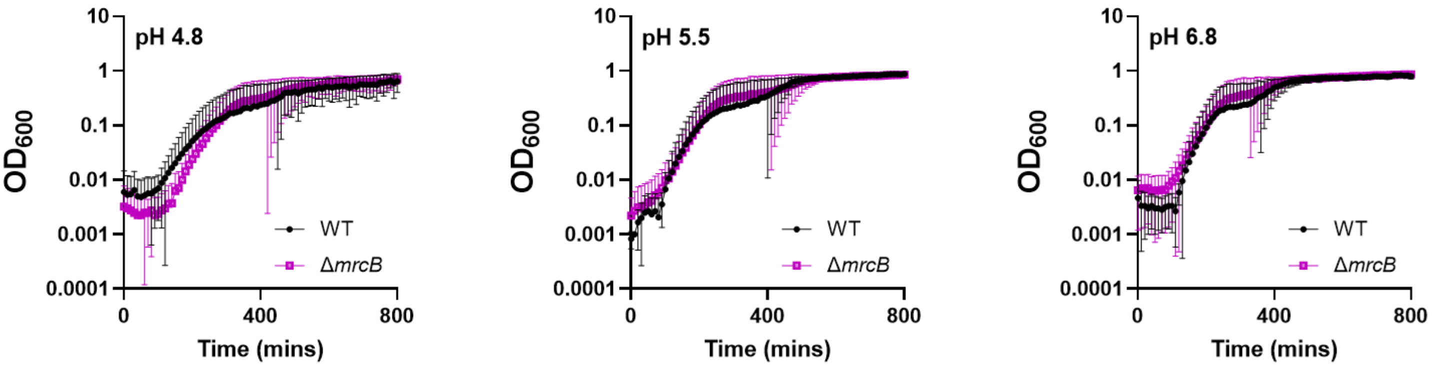

pH 4.8
pH 5.5
pH 6.8
OD600
10
1
0.1
0.01
0.001
0.0001
0
400
800
Time (mins)
WT
ΔmrcB

Sfig 6

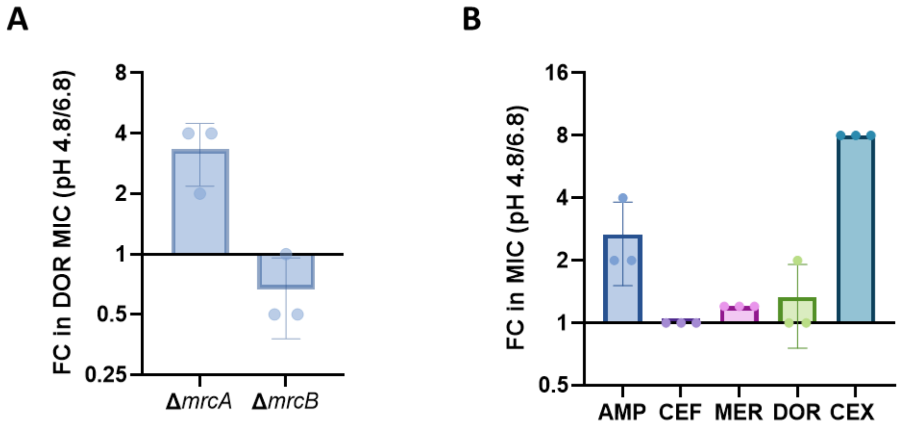
A
FC in DOR MIC (pH 4.8/6.8)
8
4
2
1
0.5
0.25
ΔmrcA
ΔmrcB
B
FC in MIC (pH 4.8/6.8)
16
8
4
2
1
0.5
AMP
CEF
MER
DOR
CEX
